# Novel microRNA multivariate biomarkers of response to immunotherapy against HPV E6 oncogene

**DOI:** 10.1101/2020.06.26.174441

**Authors:** Rebecca M. Fleeman, Gina Deiter, Kristen Lambert, Elizabeth A. Proctor, Rébécca Phaëton

## Abstract

Cervical cancer is caused by the persistent infection high-risk types of human papillomavirus (HPV) in over 99.9% of cases. To favor malignant transformation, HPV E6 and E7 oncogenes disrupt both p53 and retinoblastoma (Rb) respectively and control microRNA (miR) networks. We have previously demonstrated the therapeutic potential of anti-HPV E6 monoclonal antibodies (mAbs) in experimental models of human cervical cancer; yet the underlying mechanism remains unclear. Here, we sought to determine if anti-HPV E6 mAbs modulate the miR signatures of HPV E6 oncogenes. To this end, we performed qRT-PCR to measure the expression of thirty-four miRs and found that univariate analysis is not able to identify novel interactions characteristic of complex biological systems. Thus, we utilized partial least squares discriminant analysis (PLSDA) to identify signatures of co-varying miRs specific to mAb treatment. These miR signatures predictively discriminate between anti-HPV E6 mAb response and control mAb treatment, which may provide mechanistic insight into the action of anti-HPV E6 mAbs.

## Introduction

Cervical cancer remains a significant cause of cancer deaths worldwide, with over 528,000 women diagnosed annually.^1^ The etiologic agent of most cervical cancer is persistent infection with high-risk types of human papillomavirus (HPV), found in over 99.9% of cases.^2,3^ When diagnosed in early stages, cervical cancer is mainly cured with definitive surgery and/or combined chemotherapy and radiation therapy.^4^ In early-stages, the five-year overall survival is greater than 90%.^5^ However, in advanced or recurrent stages, where metastatic spread is beyond the cervix, treatment options are limited and the overall five-year survival is poor (5-25%).^6^

To address the urgent need for novel therapies, our laboratory previously demonstrated the significant therapeutic advantage and limited toxicity profiles of targeted therapy using monoclonal antibodies (mAbs) against human papillomavirus (HPV) E6 and E7 oncogenes. We showed a strong effect of both C1P5 (anti-HPV E6, AbCam Inc.) and TVG701Y (anti-HPV E7, AbCam Inc.) to inhibit tumor growth and activate the immune system, with limited toxicities in experimental animal models of cervical cancer.

In addition to deregulation of p53 and retinoblastoma (pRb) pathways by HPV E6 and E7 respectively, HPV-induced carcinogenesis of the cervical epithelium into invasive cervical cancer relies on both gaining transcriptional and translation control of microRNA-messenger RNA (miR-mRNA) regulatory networks.^7–10^ These small non-coding RNAs regulate over 30% of protein-coding genes, targeting several mRNAs simultaneously, immortalizing cells, and leading to malignant transformation.^11^ Moreover, HPV E6 and E7 oncogenes promote the malignant phenotype of uncontrolled cell proliferation, differentiation, and inhibition of apoptosis in part by regulating miR-mRNA networks.^12–15^

In an effort to gain further insight, we hypothesized that perhaps miRs are differentially expressed in response to treatment with mAbs. Here, we focused anti-HPV E6 oncoprotein treatment with mAb C1P5 to first confirm the capacity of the candidate miRs to distinguish between HPV statuses. We identified thirty-four candidate miRs by performing a scoping review of the literature and selected the miRs that were reported to be significant in cervical cancer and associated with HPV E6 and E7 oncogenes to identify C1P5 treatment-associated signatures unique to targeted immunotherapy.^9,11,14,16^ Our overarching goal is to uncover novel miR networks that identify novel mechanisms or have the potential to be utilized as a biomarker of therapeutic response.

Notably, in complex biological systems where several interactions and networks have constant interplay, sensitive mathematical modeling of dynamic expression is necessary for identifying subtle changes in expression with nonetheless high impact on treatment response. We utilized computational modeling with the supervised learning tool partial least squares discriminant analysis (PLSDA) to find a biomarker-relevant miR signature of C1P5 treatment. Indeed, our previous studies show that signatures of highly interactive variables can be more predictive of disease state than single entities alone.^17–20^ Thus, PLSDA was necessary to find meaningful miR signatures that are undetectable using univariate and unsupervised modeling, due to the greater sensitivity in detecting subtle and covarying changes.

Here we report novel signatures of C1P5 treatment response that can be further studied to uncover additional mechanisms, explore alternate therapeutic vulnerabilities that uncouple miR-mRNA networks, and potentially lead to highly efficacious treatments that translate to clinical use.

## Results

### A. C1P5 treatment does not have a temporal effect on miR signature

To confirm the stability of our miRs, we evaluated fold expression changes using RT-qPCR over 4- and 8-hours of treatment with C1P5 mAb. Our previous work demonstrates that in vitro treatment of cervical cancer cell lines with C1P5 mAb in cell culture causes cell death via initiation of apoptotic pathways.21–25 While the effect of C1P5 impact peaked at an eight hour time-point, in-*vivo* murine models demonstrated sustained inhibition of growth of experimentally derived cervical cancer tumors for eighteen-days. Thus we patterned our investigations of miR signatures based on knowledge gained from previous experimentation to determine if there is a time-dependent regulation of miRs.^26^

Examination of miR fold changes showed no changes in either C33A or SiHa cell lines over time. PLSDA of miR expression at 4 and 8 hours post-treatment demonstrated that miR fold change expression was indistinguishable between time points of the same treatment conditions, suggesting no time-related impact of profiles. Furthermore, analysis of individual miRs did not clearly discern HPV-positive or HPV-negative status at these time points.

### B. Univariate analysis is insufficient to detect miR expression signatures

Using a targeted screening approach of thirty-four miRs regulated by HPV E6 and E7, we compared expression profiles of the SiHa (HPV-positive) and C33A (HPV-negative) cervical cancer cell lines. We first performed real-time quantitative polymerase chain reaction (RT-qPCR) analysis of E6 and E7 mRNA in nascent cell lines and confirmed that C33A does not express HPV E6 or E7 mRNA (Figure S1). Therefore, we expected to find that the treated SiHa cells would be easily distinguishable from untreated SiHa cells, as well as from the C33A cell line, by individual E6- and E7-related miR levels. However, when we utilized univariate t-tests to compare the fold-expression changes of C1P5 and MOPC-21 treatments, we found no statistically significant changes in expression of the HPV-related miRs between treatment groups in either cell line (Figure S2).

### C. Supervised machine learning using PLSDA distinguishes miR signatures of HPV cell types

Because miR expression levels are highly interdependent, treating them as independent variables can lead to missing important information. To build a predictive miR expression signature specific to HPV E6 and E7 expressing cervical cancer, HPV positive, we performed supervised machine learning using partial least squares discriminant analysis (PLSDA).^27^ PLSDA accounts for the interdependent miR expression profiles when distinguishing groups. Indeed, our PLSDA model of SiHa vs. C33A untreated cells identified a set of infection-predictive miRs, highlighting the overwhelming difference in miR regulation with HPV status (Figure 1). Variable importance in projection (VIP) scores were quantified to assess the importance of each miR to the prediction accuracy of the model. miR-9, 15a, 16, 23b, 27a, 27b, 29a, 34a, 99a, 100, 148, 155, 199a, and 218 most significantly contributed to the distinction between SiHa and C33A cell types in our model.

**Figure 1.**
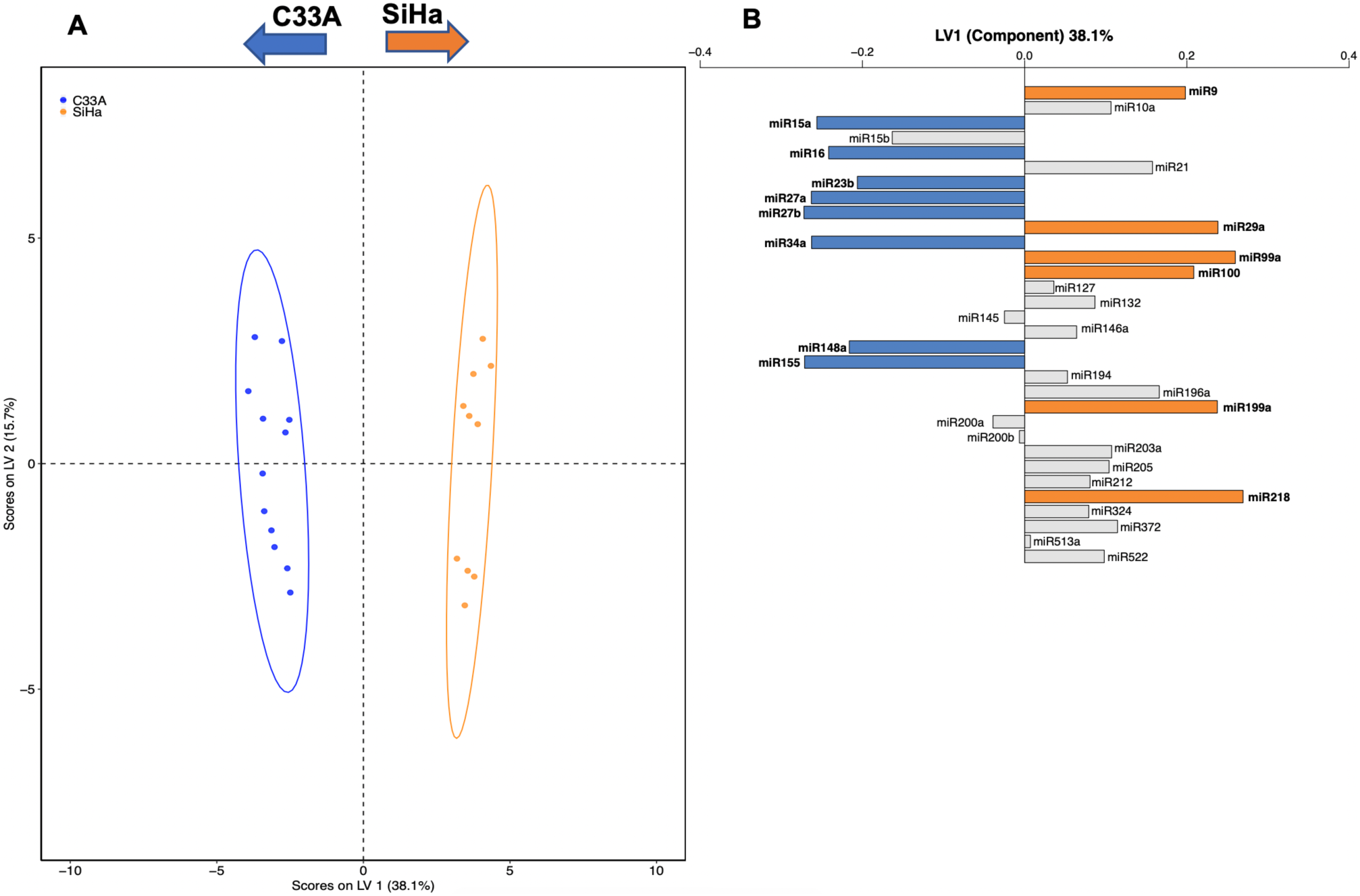
PLSDA separates miR expression by cell type. **(A)** Scores plot of PLSDA for C33A and SiHa untreated cells (2 LVs, 100% model accuracy, 100% confidence). **(B)** Latent variable (LV) 1 loadings plot for PLSDA of C33A and SiHa untreated cells. Orange and blue colored loadings have a VIP score >1.

### D. Strong miR expression signatures predictively identify C1P5 mAb treatment

We then identified a signature of miRs distinctive to mAb treatment, where variance in parameters most correlated with either C1P5 or MOPC-21 treatment were projected onto latent variable 1 (LV1). Using the VIP cytokines from initial PLSDA, we constructed a distinct signature of key miRs whose co-varying changes in expression are predictive of treatment group (Figure 2). To iterate for variable selection, we conducted a second round of PLSDA with only parameters with VIP score > 1 in the first round (Methods). When comparing miR expression changes in SiHa (HPV positive) cells treated with C1P5, an anti-HPV E6 mAb, to untreated SiHa cells, miR −99a, −127, −155, −200a, and −200b emerged as the greatest contributors to differentiating the two groups along latent variable (LV) 1 (Figure 2A). miR-127 was upregulated by C1P5 treatment, while miR −99a, −155, −200a, and −200b were down-regulated by C1P5, as evidenced by their directional contribution in the loadings on LV1 (Figure 2A). This finding indicates that decreased expression of the latter group of miRs is an important response to mAb treatment of HPV in infected cells. The resulting model for distinguishing C1P5-treated from untreated SiHa cells had an accuracy of 64.2%, as compared to random predictive ability of 47.1% (87.0% confidence). We next evaluated whether we could identify a similar pattern in C33A (HPV negative) cells. Iterating for variable selection by using two rounds of PLSDA, we again were able to identify a signature of miRs which separate C1P5 treatment from untreated cells (72.0% accuracy, 97.5% confidence) (Figure 3).

**Figure 2.**
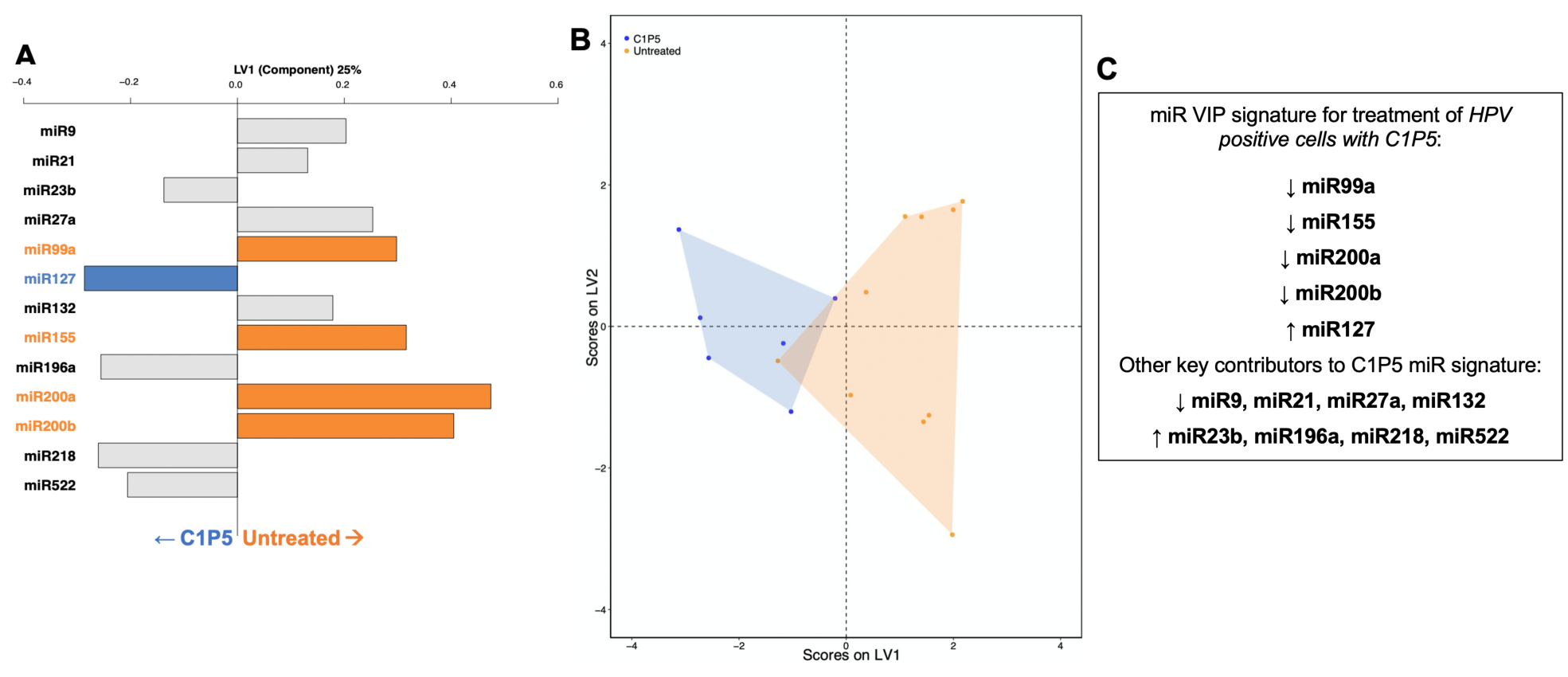
Strong miR expression signatures predictively identify C1P5 mAb treatment in SiHa cells. **(A)** PLSDA loadings and (B) scores plot for miR signatures of SiHa C1P5 vs Untreated, accuracy 64.2%, confidence 87.0%. **(C)** VIP E6/E7 relations in SiHa cells treated with C1P5.

**Figure 3.**
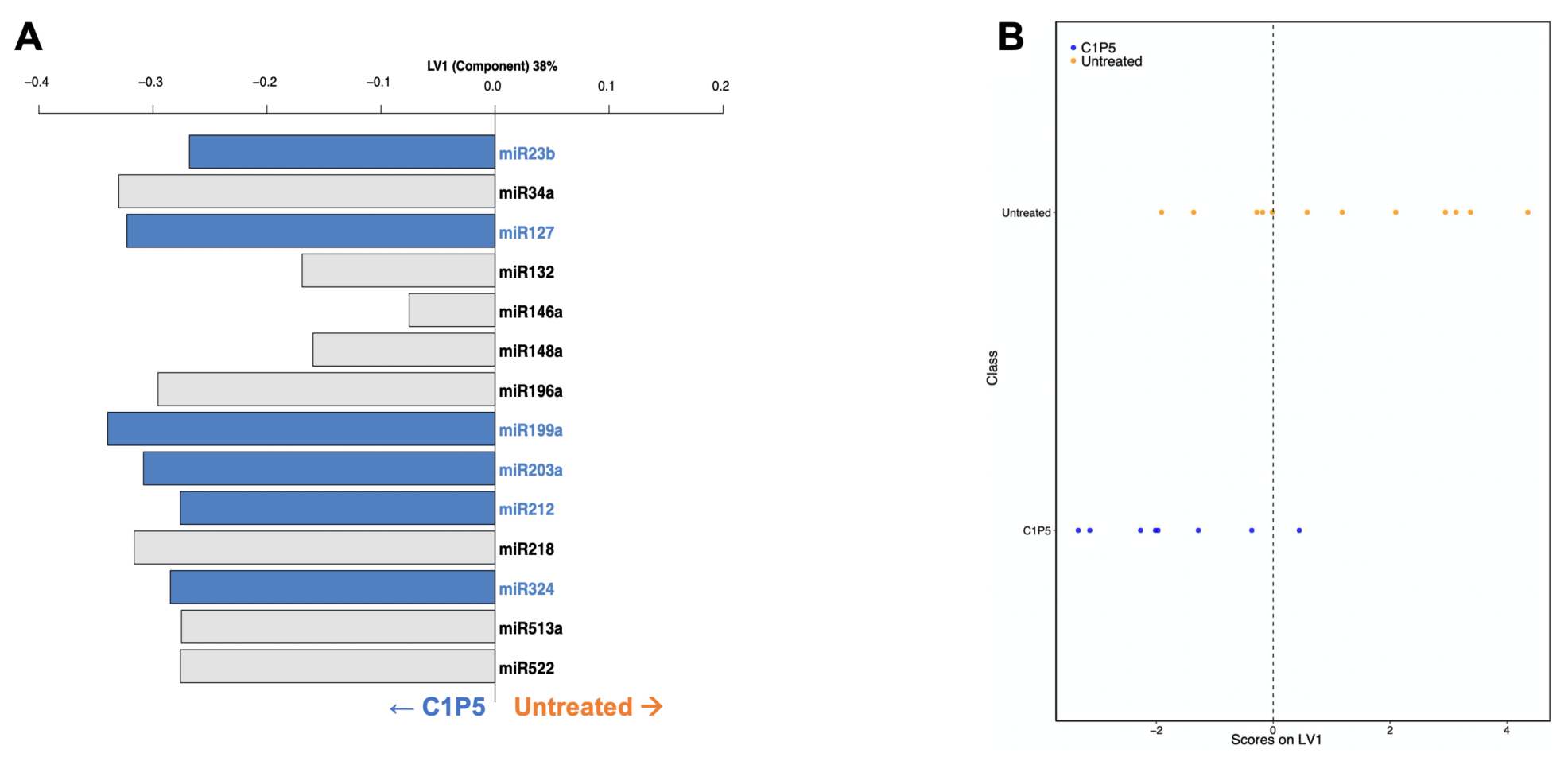
miR expression signatures predictively identify C1P5 mAb treatment in C33A cells. **(A)** Loadings plot and **(B)** scores plot for miR signatures of C33A C1P5 vs untreated, accuracy 72.0%, confidence 97.5%.

Finally, as a proof of concept, we also treated SiHa cells with MOPC-21, a non-specific IgG mAb, and compared miR expression with untreated cells, with the expectation that miR expression patterns between the two would be indistinguishable. Indeed, the resulting PLSDA model made from only parameters with VIP score > 1 in the first round had a 55.4% accuracy, compared to random predictive ability of 46.0%, and only 13.8% confidence, indicating that signatures found were no better than random modeling. Overall, these results indicate that in both HPV positive and negative cells, strong miR expression signatures can clearly delineate signatures of C1P5 anti-HPV E6 mAb treated versus untreated cells.

## Discussion

Cervical cancer remains a major cause of cancer related deaths especially in underserved minority populations.^2,28^ Carcinogenesis into invasive cervical cancer occurs in a multi-step process with the hallmarks of p53 deregulation by E6 and the disruption of retinoblastoma (rB) by E7.^7,29,30^ Moreover, the pathogenesis of HPV E6 and E7 has been shown to rely heavily on miR networks to promote the metastatic phenotype of cervical cancer.^31–34^ To this end, we evaluated 34 miRs regulated by HPV E6 and E7 oncogenes and were able to confidently detect specific differential expression profiles strictly attributable to C1P5 mAb therapy in HPV positive cells.

A growing body of literature supports the considerable diagnostic and prognostic value of miR signatures in several disease processes.^31,35–38^ Thus, miRs are rapidly becoming a common target of investigation of clinical benefit for both biomarkers and therapeutic targets. This means that targeting an oncogenic miR can serve to inhibit an entire malignant network of mRNA messages.^39^ Furthermore, the stability of miRs in biofluids in addition to their widespread bioavailability makes them attractive drug targets as well as potential biomarkers of therapeutic response.

In clinical trial design, the most common determinant of tumor responses for solid tumors are the RECIST (Response Evaluation Criteria In Solid Tumors) criteria that utilize radiographic imaging to quantify tumor burden and determine response to therapy by noting decreasing measurement of tumor size and calculations of tumor volume.^40^ However, the detection of immune-related responses require novel biomarkers to identify appropriate therapeutic endpoints of immunotherapy trials.^41–43^

To our knowledge, this work is the first analysis of miRs in response to mAb-targeted therapy in cervical cancer with anti-HPV E6 oncogene C1P5 mAb therapy. While we expect that tumor growth inhibition will occur in future Phase 1 Clinical trials of C1P5 mAbs, initial responses may be on a cellular level initially. Thus identification of a biomarker is paramount. Several authors have published miR signatures in cervical cancer in response to chemotherapy, radiation, or surgery.^44,45^ However, the lack of agreed method of RNA acquisition and the differences in statistical approaches renders diverse findings and thus minimal clinical impact.

In this report we have identified miR signatures unique to C1P5 mAb therapy. Our results indicate a specific miR expression signature corresponding to the greatest differences in mAb anti-HPV E6 treatment responses of miR expression in HPV-positive cells. We found that these subtle biological changes necessitate the use of supervised machine analysis in order to distinguish the significant changes in miR expression from biological noise. Comparison of the miR therapeutic C1P5 response signature to the non-targeted mAb MOPC-21 treatment signature allowed us to evaluate the potential contribution of mAbs to miR expression, based on their inherent immunogenicity.

While immunogenicity does result in a specific miR network signature, it is distinct from the treatment signature, suggesting that the therapeutic response to C1P5 is not purely due to immunogenicity. Our data concurs with this known fact as miRs were noted to be differentially expressed by MOPC-21. However, these were not the same miRs specifically altered by C1P5 mAb therapy. This is critical as immunogenicity can be non-specific and attributable to generalized mechanisms of interactions with mAbs.

PLSDA is an established technique. However, our use of PLSDA for miR expression changes following mAb treatment is first-of-kind. Major advantages of this supervised method over more common unsupervised multivariate dimensionality reduction methods, such as principal component analysis (PCA), include the ability to choose parameters of variance in order to answer a specific question, while retaining the interpretability of a linear modeling technique. Additionally, PLSDA allows for calculation of VIP scores, providing a basis for variable selection for a more specific signature. Of note, univariate analysis of changes in miR expression is common, yet, as we have shown here, may be misleading due to the interdependency of miR networks. Thus, supervised machine learning methods, like PLSDA, which account for the covarying mechanism of expression, should be emphasized.

In order to define the application of these miRs as either a biomarker or as a potential therapeutic target, we plan to perform pathway analysis of miR-mRNA networks to uncover their significance. The ability of our models to identify changes in HPV-relevant miR networks may be vital to early detection of HPV status, tracking progress of treatment, and identifying better treatments for diverse patients.

In our efforts to develop mAbs for therapeutic use, it is necessary to concomitantly develop a meaningful way to assess response and treatment effect. The ability of the relevant biomarkers we found here to diagnose disease or to provide an assessment of treatment efficacy and progress must be tested in the future before being used in a clinical setting. Thus, our future studies will uncover the mechanisms by which the mAb C1P5 uncouples oncogenic miR-mRNA regulated networks and if anti-tumor effects can be harnessed.

## Materials and Methods

### E. Cell lines and culture conditions

Both cervical cancer cell lines were attained from American Type Culture Collection (ATCC Manassas, VA), C33A (HPV-negative) and SiHa (HPV-positive, 1-2 viral copies/cell). C33A and SiHa cervical cancer cells were both cultured in Eagle’s Minimum Essential Medium (EMEM). EMEM was supplemented with 10% fetal bovine serum (FBS, Sigma) and 1% Penicillin (10,000 U)–Streptomycin (10 mg/ml) solution at 37°C in a 5% CO_2_ incubator. When cells reached 70% confluence, media was changed to serum-free media prior to the addition of C1P5 mAb.

### F. Antibodies

Murine mAb C1P5 (IgG1 against HPV-16 E6 + HPV 18-E6) was purchased from (Abcam Inc., Cambridge, MA) and MOPC-21 (BD Biosciences, San Jose, CA) was chosen as an IgG1 isotype murine control antibody to characterize non-specific treatment response elicited by C1P5 mAb. Phosphate buffered saline (PBS) was added to fresh media for untreated cell conditions as a vehicle.

### G. RT-qPCR

Total RNA was isolated from cell lines using TRIzol reagent (ThermoFisher Scientific, Waltham, MA) and Direct-zol RNA Miniprep Kit (Zymo Research Corp., Irvine, CA) then reverse transcribed into cDNA using qScript microRNA cDNA Synthesis Kit (Quanta Biosciences, Beverly, MA) according to the manufacturer’s instructions. Quantitative PCR was performed to determine the expression of many microRNAs using iTaq Universal SYBR Green Supermix (Bio-Rad, Hercules, CA) according to the manufacturer’s protocols. The 2^-ΔΔCt^ method was used for analysis of the microRNA qPCR data. miR 25 was utilized as endogenous control based on the stability of its expression in both cell lines. Many of the miR primers were supplied by the Faoud Ishmael lab, Hershey, PA, as well as ordered from Integrated DNA Technologies, Coralville, IA.

### H. Data Analysis

Univariate analysis was conducted in Prism 8 (v 8.4.1). Multiple t-tests were used with discovery determined using the Two-stage linear step-up procedure of Benjamini, Krieger and Yekutieli, with Q = 1%. Each miR was analyzed individually, without assuming a consistent standard deviation.

Partial least squares discriminant analysis (PLSDA) is a supervised machine learning tool for understanding interdependent changes in a set of measured variables as they relate to an outcome or grouping of choice. PLSDA uses linear combinations of variables (miR expression) to predict the variation in dependent variables (mAb treatment or cell type).^27,46^

The miR expression data sets measured at 4- and 8-hours were combined for analysis because no significant temporal differences in expression were seen between these time points. PLSDA was performed in R (v.3.6.1) using ropls (v.1.16.0)^47^ and ggplot2 (v.3.2.1)^48^ packages. Data were mean-centered and unit-variance scaled. Cross-validation was performed with one-third of the data. Model confidence was calculated by comparing model accuracy to the accuracy distribution of 100 models constructed with randomly permuted class assignment. When stated, we performed a second round of PLSDA using only those miRs with VIP score >1 in order to remove noise and improve predictability of the model.

All PLSDA models were orthogonalized on the first latent variable (LV1), so that variance in parameters most correlated with the groupings of interest was projected onto LV1. The number of latent variables was chosen to minimize cross-validation error. Variable importance in projection (VIP) scores were calculated to quantify the importance of each given parameter to the prediction accuracy of the overall model. VIP scores were calculated by averaging the weight of each miR on every latent variable across the entire model, normalized by the percent variance explained by each respective latent variable:

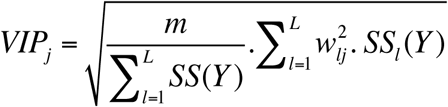

where *m* is the total number of prediectors, *l* is the latent variable, *L* is the number of latent variables in the model, *w*_*lj*_ is the weight (inverse loading) of predictor *j* on latent variable *l*, and *SS*_*j*_*(Y)* is the variation in class Y explained by latent variable l. We note that, as a consequence of normalization, the average of squared VIP scores is 1, thus VIP > 1 is our criterion for a parameter having greater than average contribution to the model.

## Acknowledgements

We thank Dr. Faoud Ishmael for sharing the miR primers and guidance on reagents and protocol We acknowledge the Institute of Personalized Medicine during the preliminary experiments and analysis of miR signatures. We thank Madison Kuhn for thoughtful input on the computational analysis. We thank the Department of Obstetrics and Gynecologic internal departmental grant funding (RP) and startup funds from the Departments of Neurosurgery and Pharmacology (EAP) for support to perform the studies.

## Author contributions

Conceptualization of utilization of miRs as a biomarker of HPV related immunotherapy RP. Data analysis EAP and RMF. Experimentation with RT-qPCR data and calculations RP and GD. Funding RP and EAP. Writing, reviewing, and editing RMF, EAP, GD and RP. Laboratory Resources RP.

