## Supplemental Figures for "Novel microRNA multivariate biomarkers of response to immunotherapy against HPV E6 oncogene"

**S1. C33A does not express HPV E6 or E7 mRNA.** Univariate analysis of RT-qPCR data for E6/E7 mRNA CasKi and SiHa, HPV (+), with E6/E7. C33A (HPV -) no HPV E6/E7.

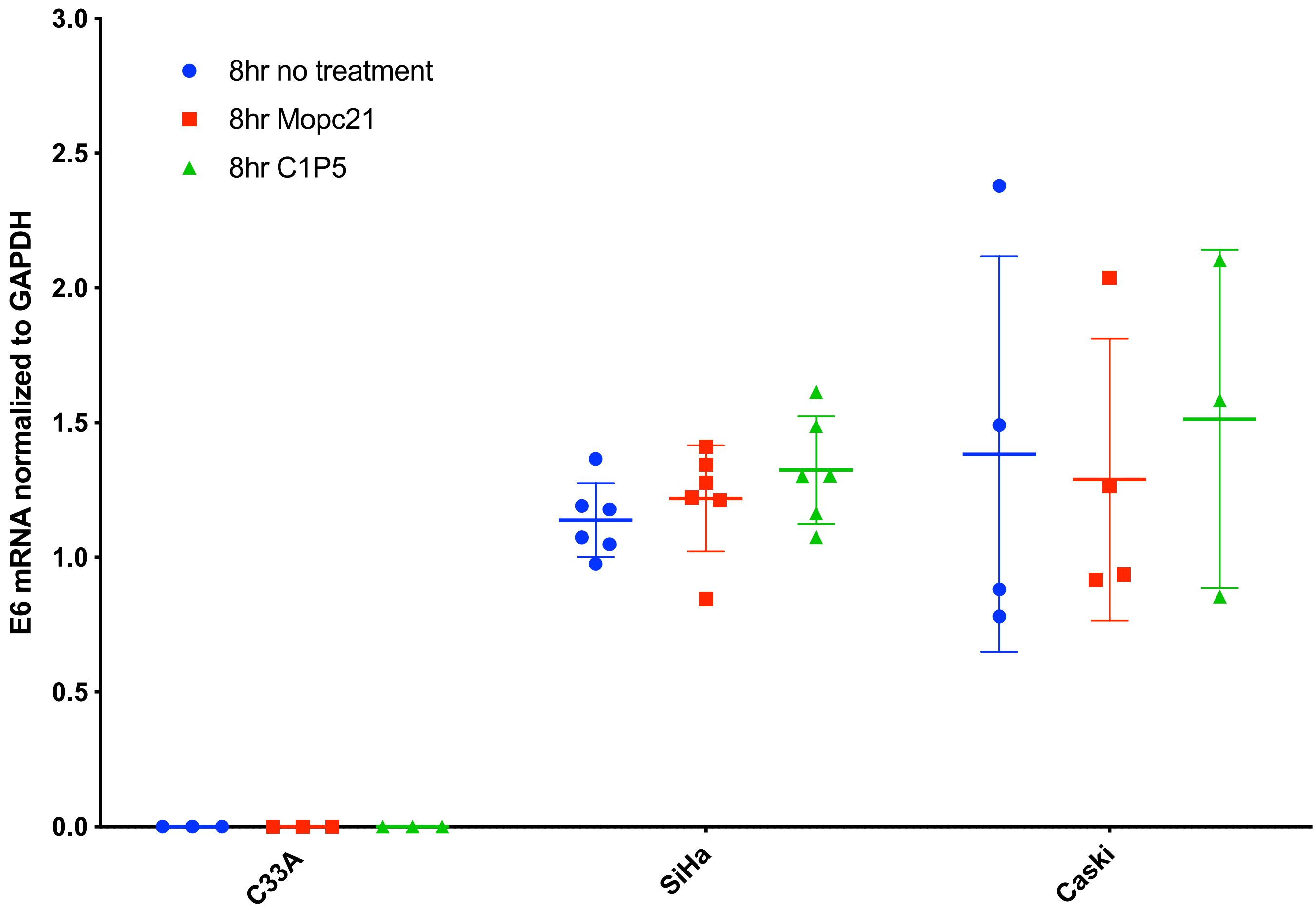

**S2. Univariate analysis is insufficient to detect miR expression signatures.** Univariate analysis of combined 4 and 8 hour miR fold expression changes measured by RT-qPCR following treatments in **(A)** SiHa HPV+ cells and **(B)** C33A HPV- cells. All comparisons measured with t-test and not significant.

**A**

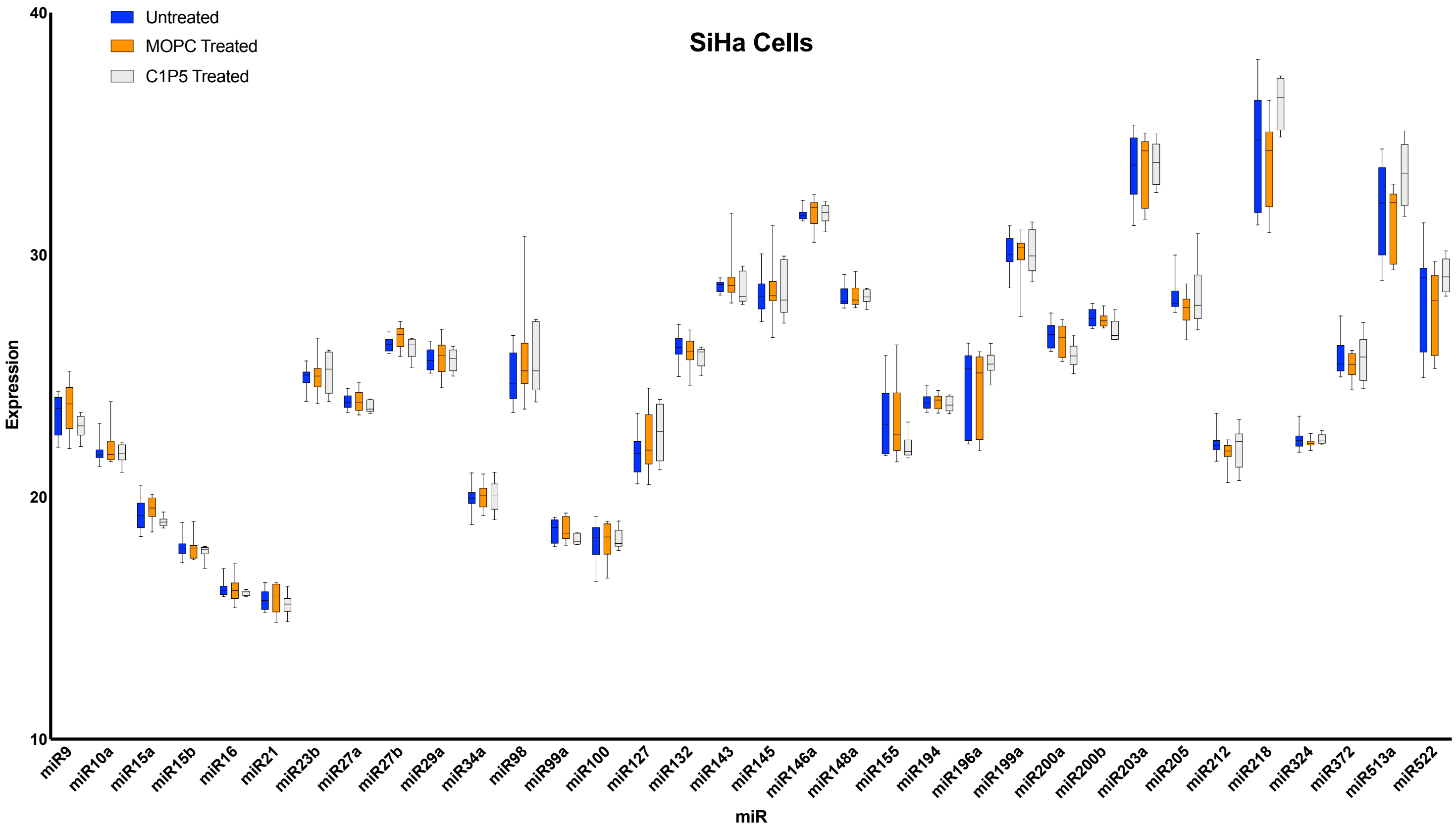


**B**

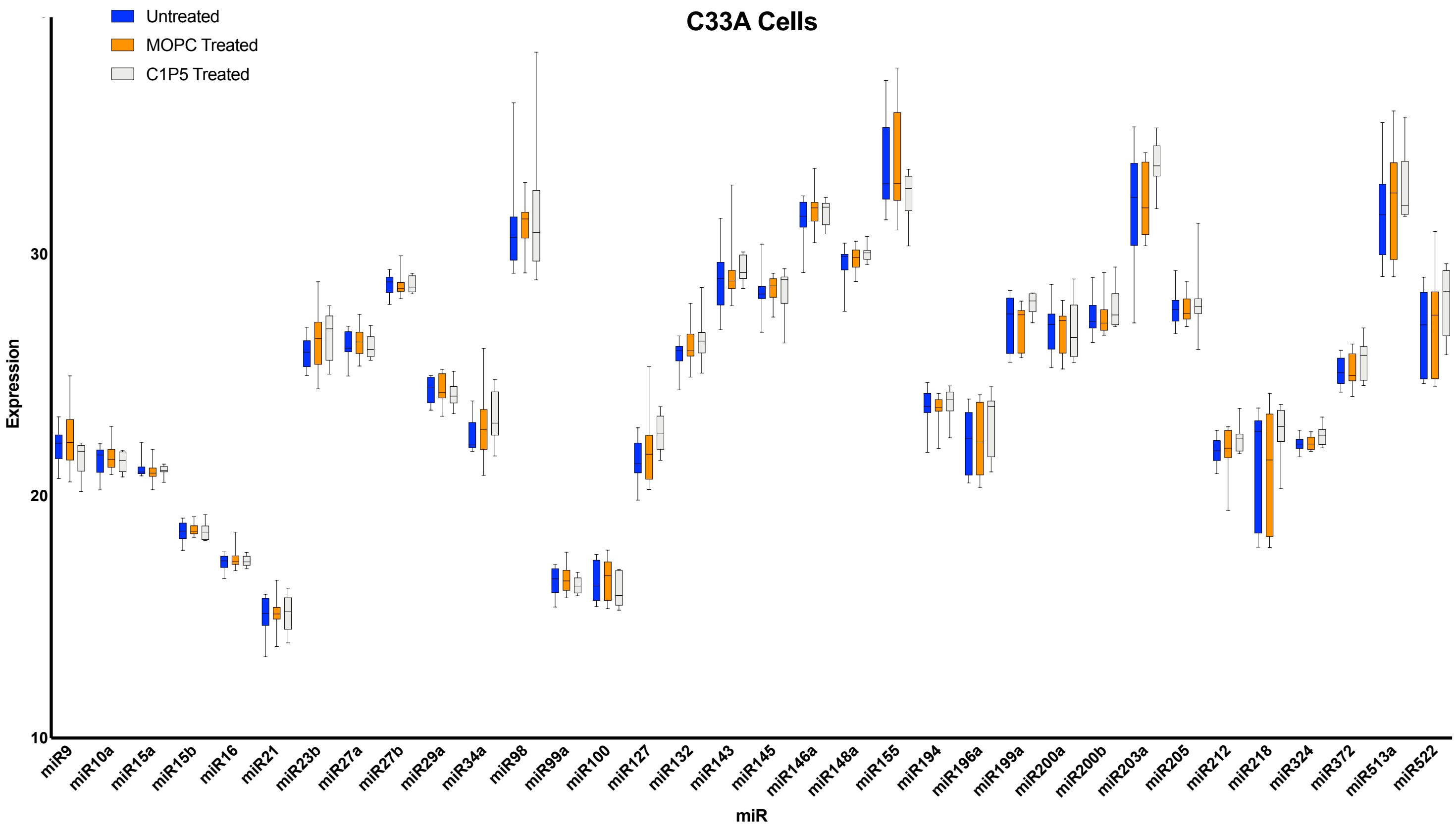
